# Inhibitors of lysosomal function or serum starvation in control or LAMP2 deficient cells do not modify the cellular levels of Parkinson disease-associated DJ-1/PARK 7 protein

**DOI:** 10.1101/246314

**Authors:** Raúl Sánchez-Lanzas, José G. Castaño

## Abstract

Mutations in PARK7/DJ-1 gene are associated with familial autosomal recessive Parkinson disease. Recently, lysosomes and chaperone mediated autophagy (CMA) has been reported to participate in the degradation of DJ-1/PARK7 protein. Lamp-2A isoform is considered as the lysosomal receptor for the uptake of proteins being degraded by the CMA pathway. We have used several cell lines with disrupted LAMP2 gene expression and their respective control cells to test the possible role of lysosomal degradation and in particular CMA in DJ-1 /PARK7 degradation. Interruption of LAMP-2 expression did not result in an increase of the steady-state protein levels of DJ-1 /PARK7, as it would have been expected. Furthermore, no change in DJ-1 /PARK7 protein levels were observed upon inhibition of lysosomal function with NH_4_Cl or NH_4_Cl plus leupeptin, or after activation of CMA by serum starvation for 24h. Accordingly, we have not found any evidence that DJ-1 /PARK7 protein levels are regulated via lysosomal degradation or the CMA pathway.

## Introduction

PARK7 /DJ-1 gene mutations are linked to autosomal recessive and early-onset clinical manifestations of Parkinson's disease. Pathogenic mutations identified in PARK7 /DJ-1 gene include CNVs (exonic deletions and truncations), and numerous missense mutations [1] [2].

DJ-1 is a dimeric protein with a flavodoxin-like structure [3–5] [6,7] and ubiquitously expressed [8]. DJ-1 wild type protein is a rather stable protein since DJ-1 protein levels remain essentially unchanged after 24h of incubation of cells with cycloheximide [9] [10]. In addition, pulse-chase experiments did not reveal a clear decay in wild type DJ-1 levels within 24 hrs [11]. Surprisingly, Wang et al [12] recently reported that DJ-1 protein levels increase after treatment of SN4741 cells with NH_4_Cl or NH_4_Cl and leupeptin for 18h. Furthermore, they show that >80% of DJ-1 is degraded after prolonged (24 hrs) serum starvation, a treatment known to activate the pathway of chaperone mediated autophagy (CMA). In CMA, proteins with a conserved aminoacid motif selectively bind to the C-terminus of the integral lysosomal membrane protein Lamp-2A, one of the three isoforms generated from LAMP2 gene transcripts by alternative splicing. After binding to Lamp-2A, proteins are translocated to the lysosome matrix for degradation [13]. The observed DJ-1 degradation upon serum starvation was prevented by co-treatment with leupeptin and NH_4_Cl, and not prevented by addition of·3-methyl adenine (3-MA). Those basic results and other experiments lead the authors [12] to the conclusion that DJ-1 protein is degraded by CMA pathway through binding to Lamp-2A [13]. Those results prompt us to replicate and validate the implication of the CMA pathway in degradation of wild type DJ-1. Using different cell lines with lack of LAMP-2 gene expression and thereby stimulation of CMA, we found no experimental evidence that CMA pathway is involved in the turnover of the DJ-1 wild type protein.

## Materials and Methods

### Ethical statement

Mice for isolation of mouse embryonic fibroblasts were handle following the ethical standards set by the National Animal Care Committee of Germany and protocol approval through the Ministerium für Energiewende, landwirtschaft und ländliche Räume, Schleswig-Holstein (V312-72241.121-3) by Dr. Paul Saftig group at the animal facilities of Institute of Biochemistry, Christian-Albrechts-Universität zu Kiel, Kiel, Germany. Human cell lines used in this study were described previously with identified source and approval for the B-cell line derived from a Danon patient [14]

### Cell lines

Human B-lymphoblastoid cell lines (B-LCL) were grown in RPMI medium (Gibco BRL) supplemented with 10% fetal bovine serum (FBS), penicillin/streptomycin and 2 mM glutamine, as described [14]. HeLa, HEK, SN4741 cells (provided by Dr. José Luis Zugaza, Faculty of Science and Technology, University of the Basque Country, UPV/EHU, Bilbao, Spain) and the different control and Lamp-2 deficient cell lines were grown in Dulbecco’s modified Eagle’s medium (DMEM, Gibco BRL) supplemented with 10% foetal bovine serum (Sigma-Aldrich) and 100 µg/mL gentamycin.. Mouse embryonic fibroblasts (MEF) were obtained from wild type (Wt) mice and MEF Lamp-2^-/y^ from Lamp-2 deficient mice [15] [16]. N2A cells transfected with shRNA scrambled sequence (shRNA scrmbl) and shRNA for Lamp-2 have been previously described [17] and were cultured in the presence of the selecting antibiotic G418 at 400 µg/mL. All cells were grown at 37ºC (except SN4741 that were cultured at 33ºC) and humidified 5% CO2. Both MEFs and N2a cell line were provided by Drs. Judith Blanz and Paul Saftig from Institute of Biochemistry, Christian-Albrechts-Universität zu Kiel, Kiel, Germany.

### Antibodies

Anti-Lamp-2A-specific antibody (Abcam ab18528) was used at 1/1000 and rabbit anti-Lamp-2A [17] was also used at 1/1000. Anti-LC-3 (Sigma) was used at 1/1000 and anti-DJ-1 (Abcam ab18257) was used at 1/2000. As control for total protein loading, antibodies against α-tubulin (1/10,000, Sigma, clone DM1a) were used.

### Studies of protein degradation

Exponentially growing cells were treated with 25 μg/ml of cycloheximide (CHX) for the times indicated. Cell viability by trypan blue exclusion was ≥95%. Cells were collected and processed for analysis by Western and immunoblot with anti-DJ-1specific antibodies. To study the basal activity of the autophagic pathway, cells in complete medium with serum were untreated (controls) or treated with 20 mM NH_4_Cl or 20 mM NH_4_Cl in combination with 50 μM leupeptin (Leup) for 24 h and processed for immunoblot analysis with anti-DJ-1 or anti-LC3 antibodies. To study the effect of serum starvation on protein levels, exponentially growing cells were washed three times with HBSS with calcium and magnesium (55037C, Sigma) and incubated in serum free medium (serum starvation) for the times indicated. Cells were then cultured in the absence or in the presence of 20 mM NH_4_Cl, 20 mM NH_4_Cl and 50 μM Leup or 10 mM 3-MA. Afterwards cells were washed three times with cold PBS and lysed in SDS-Laemmli loading buffer and processed for Western and immunoblot analysis (see below).

### Immunoblot analysis

Cells were lysed in SDS-Laemmli loading buffer, boiled for 5 min and loaded onto 10–14% SDS-PAGE (as required) and transfer to PVDF. Membranes were incubated with the corresponding primary antibodies (as indicated) and developed with anti-rabbit or anti-mouse peroxidase-labeled antibodies (1/5,000, Bio-Rad). Blots were imaged with a chemiluminiscent detector (MFChemiBIS 3.2, DNR Bio-Imaging Systems) and analyzed by quantitative densitometry using Totallab TL100 software. Protein levels were normalized respect to tubulin (protein loading control) and are expressed as mean ± s. e. m. from three different experiments.

## Results

### DJ-1 steady-state protein levels in control and LAMP2 interrupted cell lines

If Lamp-2A is implicated in the degradation of DJ-1 by the CMA pathway as reported [12], it would be expected that the interruption of LAMP2 gene expression would lead to an increase in DJ-1 protein levels. To test the validity of this prediction DJ-1 protein levels were measured in mouse embryonic fibroblasts (MEF) derived from control and LAMP2 KO mice [16], as well as in control (scrambled shRNA) and LAMP2 –deficient (shRNA for LAMP2) N2a cells obtained by stable transfection of scrambled and LAMP-2 shRNA [17]. Lack of LAMP-2(A) expression in these cell lines has been validated by using LAMP-2 and LAMP-2A specific antibodies [17], Disruption of LAMP-2 expression does not affect Lamp-1 protein expression, another abundant integral membrane lysosomal protein (see Supplementary Fig.1). We also used B-lymphoblastoid cell lines (B-LCL) derived from a control and Danon disease’s patient [14], an X-linked human disease caused by mutations in LAMP-2 gene.[18]. The data presented in Fig. 1 show that there was not significant change in DJ-1 protein levels compared to the corresponding controls in LAMP-2-deficient MEFs, N2a cells and B-LCL derived from a Danon disease patient. These results reveal that DJ-1 steady state protein levels are not significantly affected by lack of expression of Lamp-2.

**Legend to Fig. 1.**
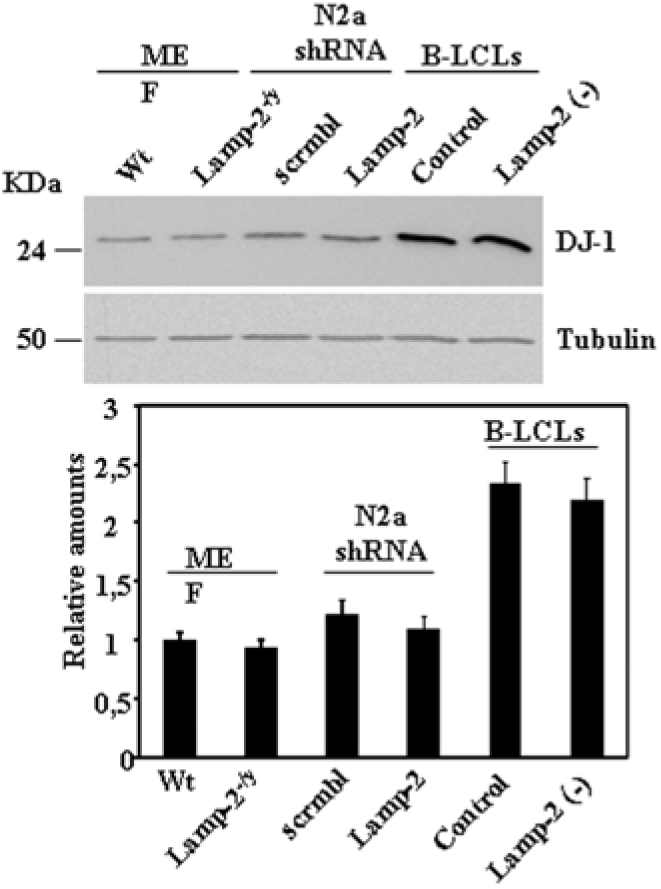
Steady-state protein expression levels of DJ-1 in control and Lamp-2-deficient cell lines. Total lysates from exponentially growing cells under basal culture conditions, control and Lamp-2-deficient MEF, N2a and B-LCLs cells were prepared and the levels of DJ-1 protein analysed by Western and immunoblot. Anti-tubulin antibodies were used as total protein loading control. Below is shown the graph of quantification of the corresponding immunoblots. Data are mean ± s. e. m. from at least three experiments. No significant difference in DJ-1 protein levels was found between controls and their corresponding Lamp-2 deficient cell lines.

### Kinetic studies of DJ-1 stability after protein synthesis inhibition

The results shown above led us to test the stability of DJ-1 in the cell lines described above after protein synthesis inhibition with cycloheximide (CHX). As we have previously described with N2a cells [10], treatment of the different cell lines under study with CHX up to 24h did not significantly change DJ-1 protein levels (Fig. 2), indicating that the half-life of the DJ-1 protein is longer than 24 hrs. Similar results were obtained (Supplementary Fig. 2) with other cell lines: HeLa, HEK293 and SN4741, the cell line used by Wang et al [12]. Under the same experimental conditions IKappaBα, used as a control, was degraded (data not shown)

**Legend to Fig. 2.**
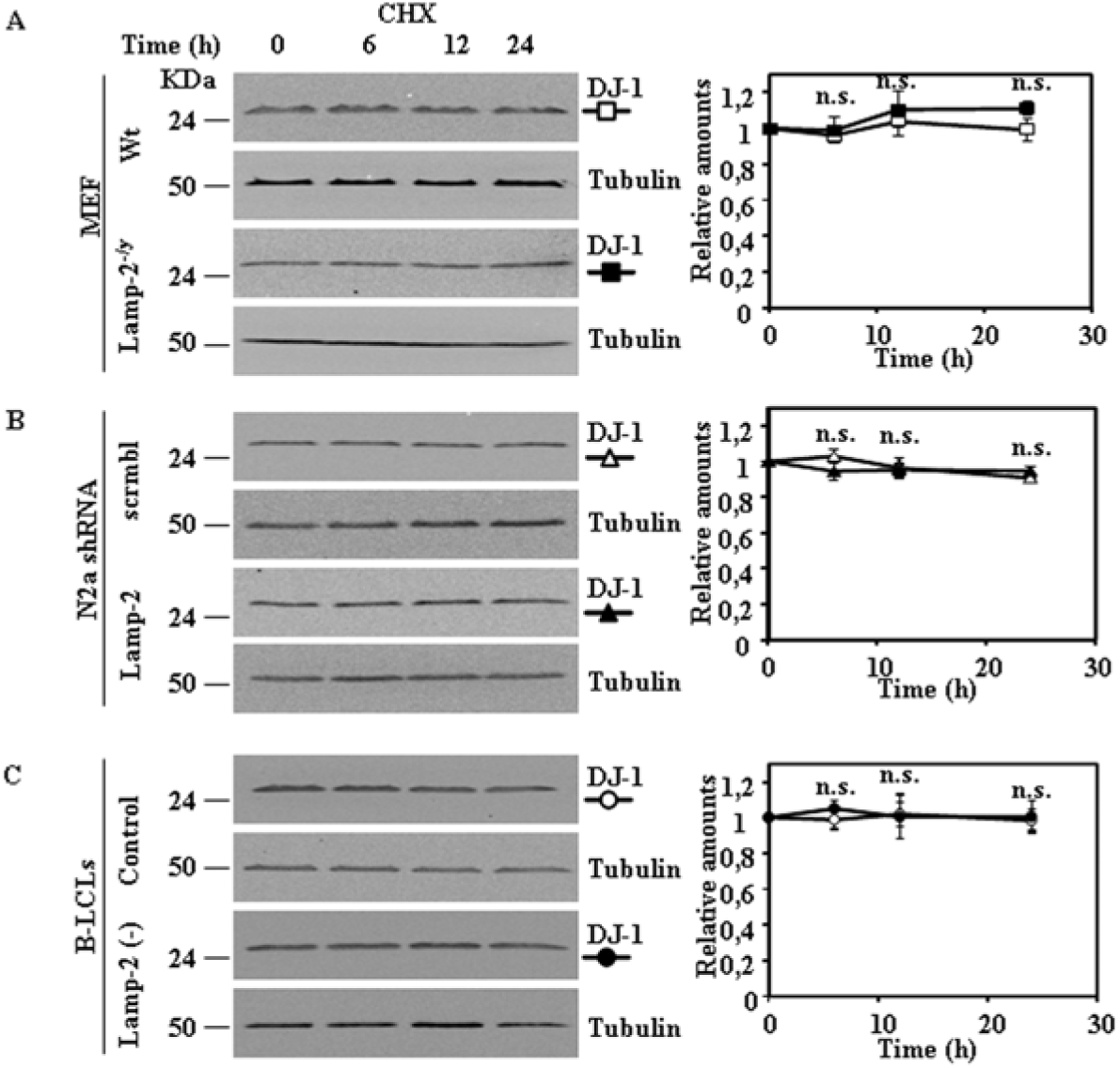
Stability of DJ-1 protein in control and Lamp-2-deficient cell lines after inhibition of protein synthesis. Exponentially growing cells from control and Lamp-2-deficient cells were treated with cycloheximide (CHX) for the times indicated. Total cell lysates were prepared and DJ-1 protein levels were analysed by Western and immunoblot with specific antibodies. Anti-tubulin antibodies were used as total protein loading control. Panels show the results obtained for MEF (A), N2a (B) and B-LCLs (C), Graphs on the right show the quantification of the levels of DJ-1 protein respect to untreated cells as controls (time 0 h). Values are expressed as mean ± s.e.m. from three different experiments. n.s. not significant difference

### Effect of lysosomal inhibition on DJ-1 protein levels

Wang et al [12] reported that inhibition of lysosomal degradation by NH_4_Cl/leupeptin treatment (18hrs) leads to increased DJ-1 protein levels. This observation led the authors to conclude that DJ-1 is degraded in lysosomes. We aimed to reproduce those results in the same cell line (SN4741) used in this study as well as in other cell lines such as MEF obtained from wild type or Lamp-2-deficient mice, N2a cells either stably transfected with scrambled shRNA or LAMP-2-specific shRNA and in B-LCLs from a control or Danon disease’s patient As shown in Fig. 3, our data showed that after 24h of treatment with NH_4_Cl or NH_4_Cl/leupeptin, there is no significant change in DJ-1 protein levels in any of the cell lines studied, including SN4741 cells. In contrast and as expected, NH_4_Cl and NH_4_Cl/leupeptin treatments increased the levels of LC3-II, as expected due to autophagic inhibition (Fig. 3).

**Legend to Fig. 3.**
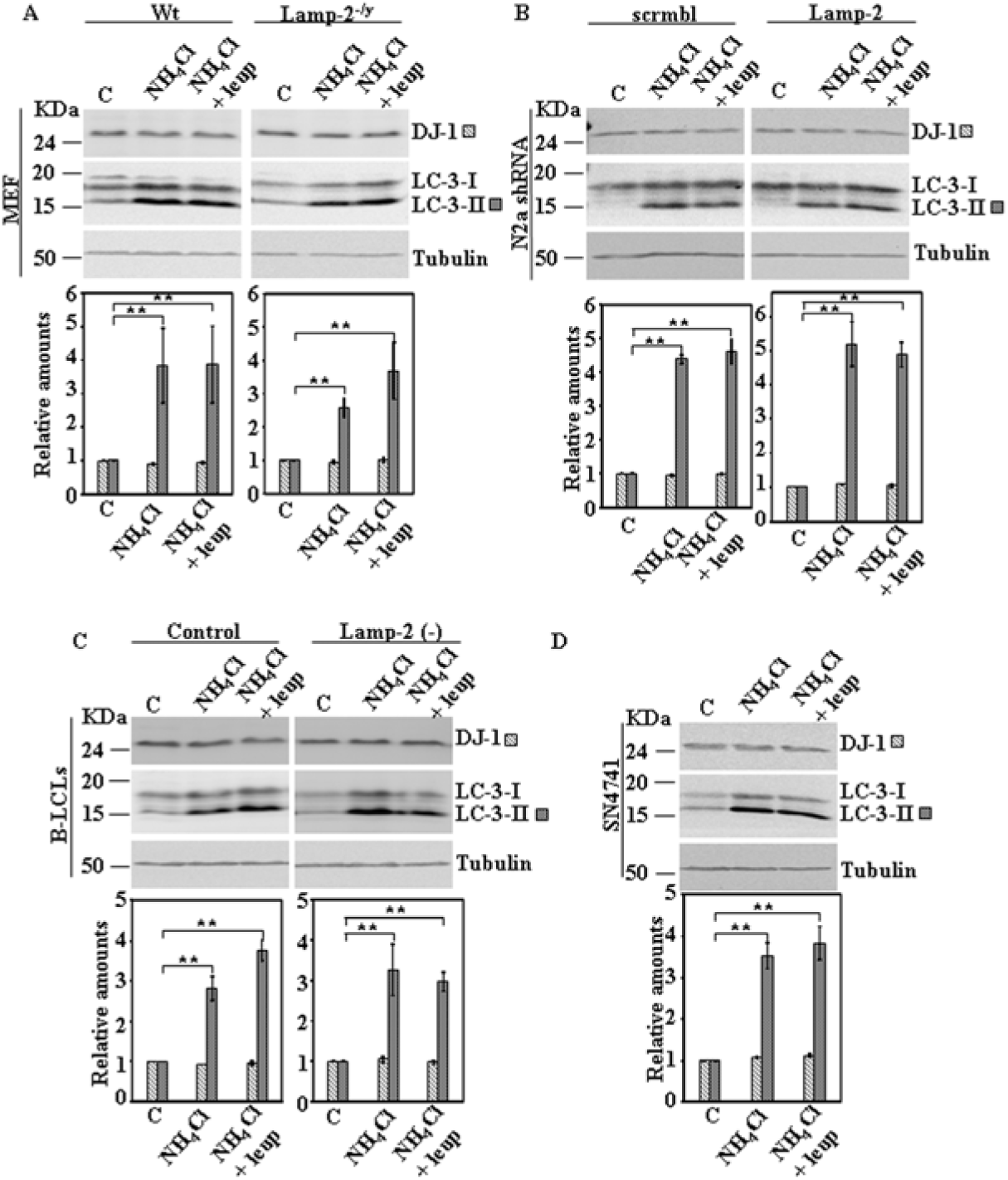
Protein expression levels of DJ-1 in control and Lamp-2-deficient cell lines in the presence of inhibitors of lysosomal function. Exponentially growing cells from control and Lamp-2-deficient cells were incubated in complete medium r suplemented with 20mM NH_4_Cl or 20 mM NH_4_Cl in combination with 50 µM leupeptin (Leup) for 24 h. Total cell lysates were analysed by Western and immunoblot with the corresponding specific antibodies: DJ-1 and LC-3. Anti-tubulin antibodies were used as total protein loading control. Panels show the results for MEF (A), N2a (B), B-LCLs (C) and SN4741 (D). Graphs below each panel show the quantification of the levels of the proteins analysed respect to the levels in cells kept in complete growth medium, controls. Values are expressed as mean ± s.e.m. from three different experiments. Significant differences between groups ** at p<0.01 by Student t-test are indicated.

### Effect of CMA activation on DJ-1 protein levels

After obtaining the above results we activated CMA by serum starvation to study its effect on DJ-1 protein levels. Serum starvation for 24h did not produce any change in DJ-1 protein levels in MEF obtained from wild type or Lamp-2-deficient mice (Fig. 4). Similar results were obtained in N2a cells either stably transfected with scrambled shRNA or LAMP-2-specific shRNA (Fig. 5). Also no significant changes in DJ-1 protein levels were found in B-LCLs from a control or Danon disease’s patient (Fig. 6) [14], when starved for 8h. In these cell lines we could not apply 24h of serum starvation since it resulted in >60% cell death, as reported previously [14]. Finally, upon CMA activation we also did not observe any change of DJ-1 protein levels in SN4741 cells (Fig. 7). Furthermore, also the addition of neither NH_4_Cl, NH_4_Cl/leupeptin nor 3-MA during the starvation period modified DJ-1 protein levels in any of the cell lines studied (see Figs. 4, 5, 6 and 7). Certainly those co-treatments during starvation resulted in the accumulation of LC3-II (NH_4_Cl, NH_4_Cl/leupeptin) and its inhibition in the presence of 3-MA (see also Figs. 4, 5, 6 and 7). Under the same experimental conditions IKappaBα, used as a control, was degraded (serum starvation) being prevented by co-treatment with NH_4_Cl, NH_4_Cl/leupeptin (data not shown).

**Legend to Fig. 4.**
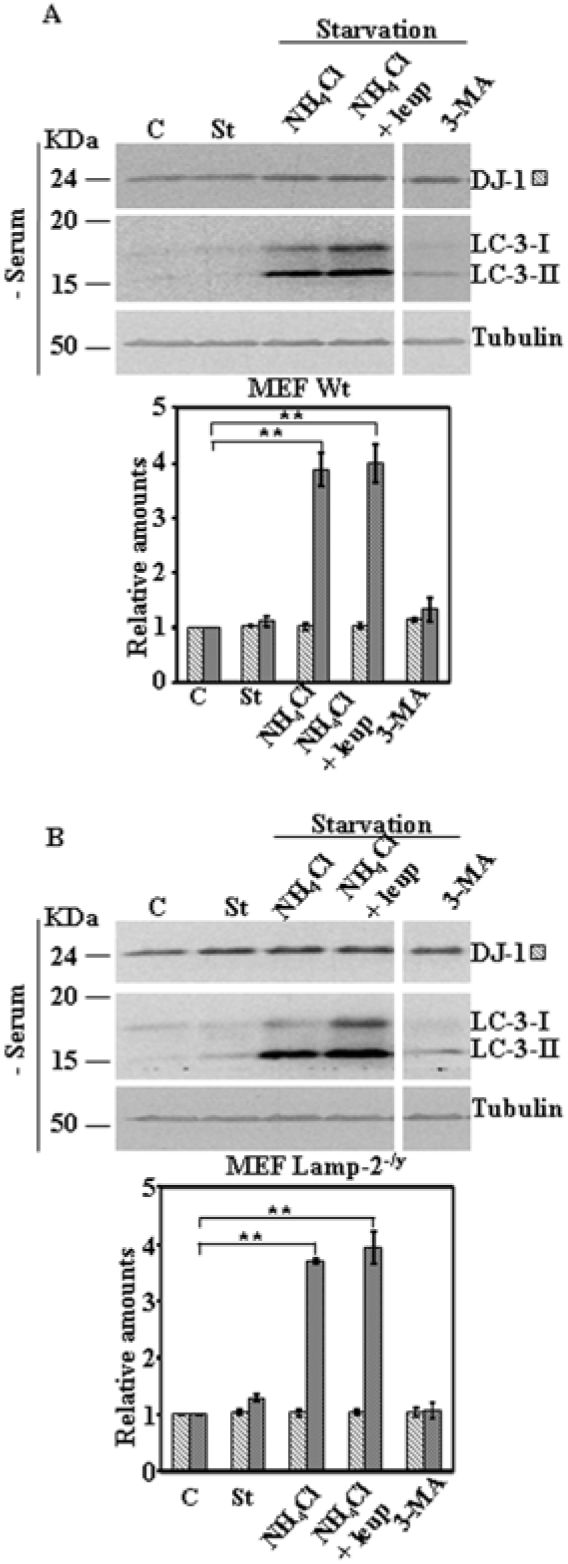
Protein expression levels of DJ-1 following activation of autophagy by serum starvation in control and Lamp-2-deficient MEF cells. Exponentially growing control and Lamp-2-deficient MEF cells were kept in complete medium (C) or starved of serum for 24 h in the absence (St) or in the presence of NH_4_Cl,NH_4_Cl and leupeptin (Leup), or 3-methyl adenine (3-MA). Panels A and B show the effect of serum starvation in MEF Wt cells and Lamp-2-deficient MEF (Lamp-2^-/y^) cells, respectively. Total cell lysates were analysed by Western and immunoblot with the corresponding specific antibodies: as indicated. Anti-tubulin antibodies were used as total protein loading control. Graphs below each panel show the quantification of the levels of the different proteins analysed respect to the levels in cells kept in complete growth medium, controls. Values are expressed as mean ± s.e.m. from three different experiments. Significant differences between groups ** at p<0.01 by Student t-test are indicated.

**Legend to Fig. 5.**
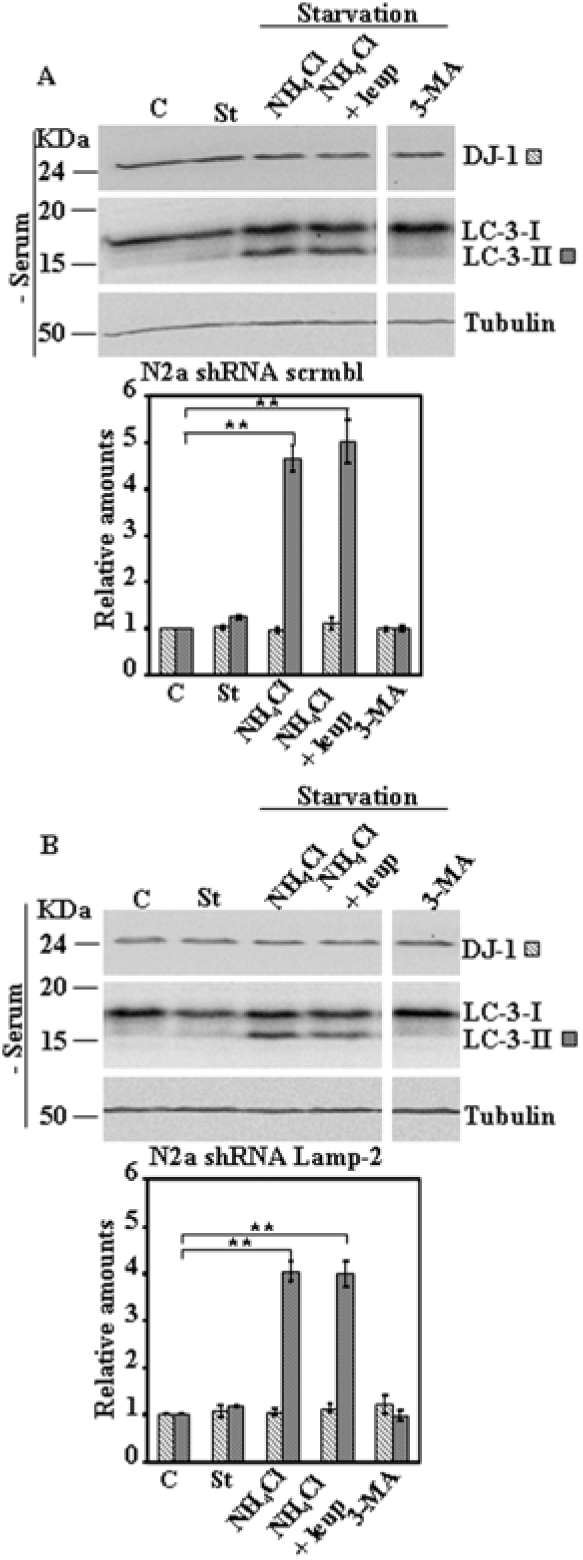
Protein expression levels of DJ-1 following activation of CMA by serum starvation in control and Lamp-2-deficient N2a cells. Exponentially growing control and Lamp-2-deficient N2a cells were kept in complete medium (C) or starved of serum for 24 h in the absence (St) or in the presence of NH_4_Cl, NH_4_Cl and leupeptin (Leup), or 3-methyl adenine (3-MA). Panels A and B show the effect of serum deprivation in N2a shRNA scrmbl cells and in Lamp-2-deficient N2a shRNA Lamp-2 cells, respectively. Total cell lysates were analysed by Western and immunoblot with the corresponding specific antibodies. Anti-tubulin antibodies were used as total protein loading control. Graphs below each panel show the quantification of the levels of the different proteins analysed respect to the levels in cells kept in complete growth medium, controls. Values are expressed as mean ± s.e.m. from three different experiments. Significant differences between groups ** at p<0.01 by Student t-test are indicated.

**Legend to Fig. 6.**
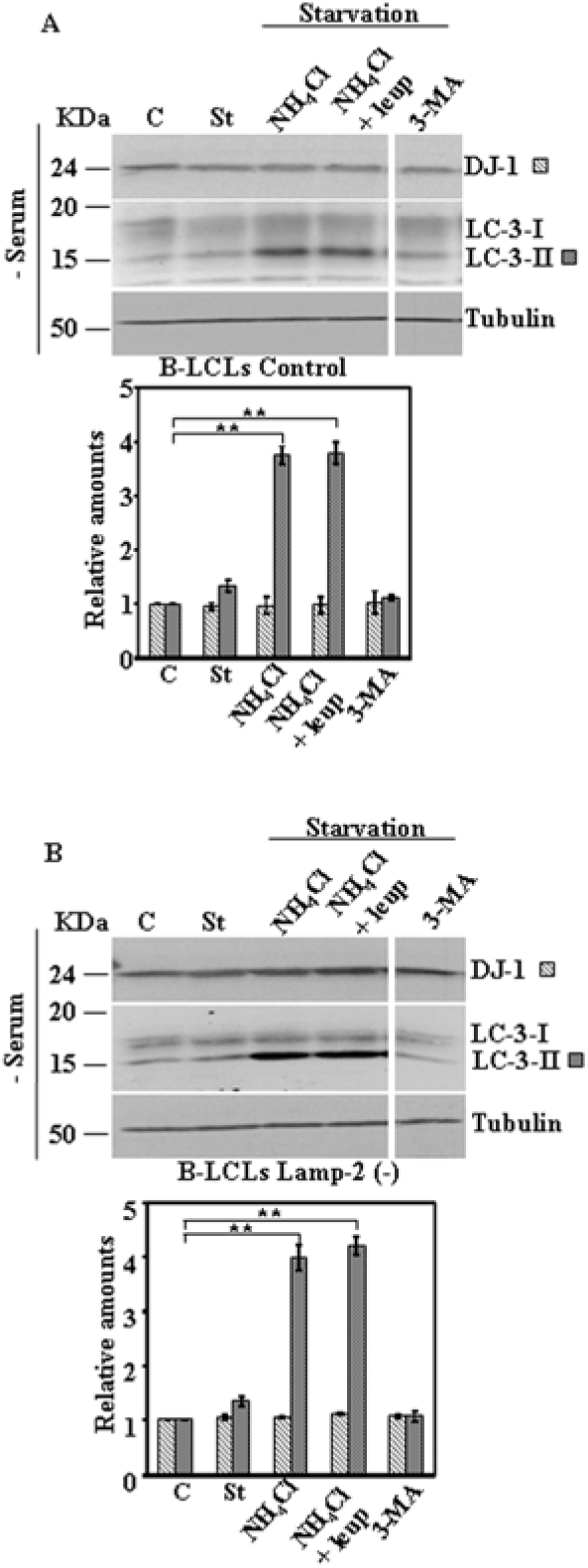
Protein expression levels of DJ-1 following activation of CMA by serum in control and Lamp-2-deficient B-LCLs. Exponentially growing control and Lamp-2-deficient B-LCL were kept in complete medium (C) or starved of serum (8h) in the absence (St) or in the presence of NH_4_Cl, NH_4_Cl and leupeptin (Leup), or 3-methyl adenine (3-MA). Panels A and B show the effect of serum deprivation in B-LCLs control and Lamp-2 (-). Lamp-2-deficient B-LCL, respectively. Total cell lysates were analysed by Western and immunoblot with the corresponding specific antibodies, as indicated. Anti-tubulin antibodies were used as total protein loading control. Graphs below each panel show the quantification of the levels of the different proteins analysed respect to the levels in cells kept in complete growth medium, controls. Values are expressed as mean ± s.e.m. from three different experiments. Significant differences between groups ** at p<0.01 by Student t-test are indicated.

**Legend to Fig. 7.**
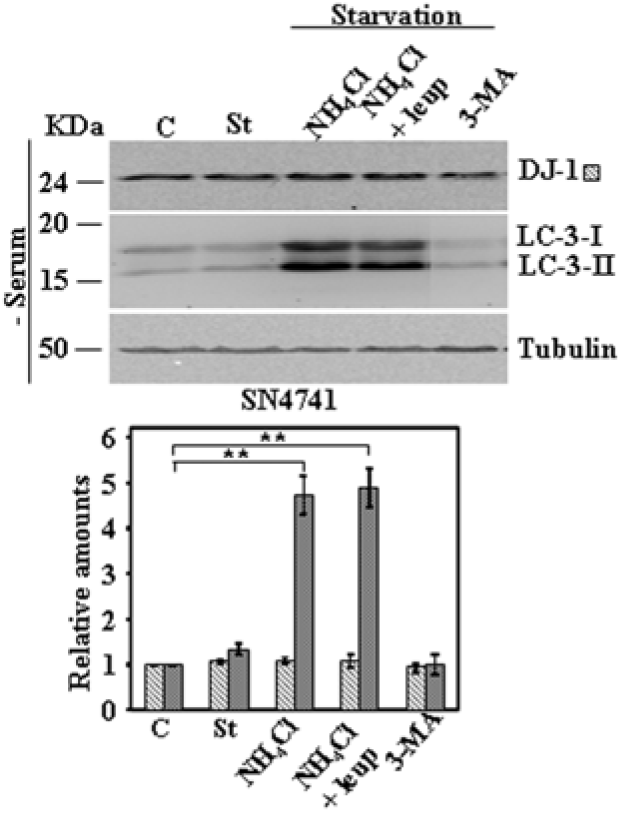
Protein expression levels of DJ-1 following activation of CMA by serum in SN4741 cells. Exponentially growing SN4741 cells were kept in completemedium (C) or starved of serum for 24 h in the absence (St) or in the presence of NH_4_Cl, NH_4_Cl and leupeptin (Leup), or 3-methyl adenine (3-MA). Panels show the effect of serum deprivation in SN4741. Total cell lysates were analysed by Western and immunoblot with the corresponding specific antibodies, as indicated. Anti-tubulin antibodies were used as total protein loading control. Graphs below each panel show the quantification of the levels of the different proteins analysed respect to the levels in cells kept in complete growth medium, controls. Values are expressed as mean ± s.e.m. from three different experiments. Significant differences between groups ** at p<0.01 by Student t-test are indicated.

### Discussion

The results presented here reveal that DJ-1 protein levels did not change in response to LAMP-2 gene expression silencing (Fig. 1) in different cell types (MEF, N2a and B-LCL), as a consequence the increased protein levels of DJ-1 reported by Wang et al [12] in SN4741 cells upon silencing LAMP-2 gene expression can not be generalized to other cell types.

DJ-1 protein is a stable protein in the cell, here we have extended previous results to several cell lines, showing that treatment with CHX for 24h did not change the amount of DJ-1 protein in MEF from wild type or LAMP-2 knock-out mice, control N2a cells and interrupted with an shRNA for LAMP-2 or in B-LCL from control or from a patient with Danon disease. Similarly, no changes were observed in HeLa, HEK293 and SN4741 cell line (Supplementary Fig. 1) used by Wang et al [12].

Treatment of cells with NH_4_Cl or NH_4_Cl/leupeptin resulted in no change in DJ-1 protein levels either in control cells or in LAMP-2 interrupted cells mentioned above (Fig. 3) and also no change was observed in SN4741 (Fig. 3D). Furthermore, the results presented here show no change in DJ-1 protein levels during starvation of serum for 24h, either in control or LAMP-2 interrupted cells (Figs. 4, 5 and 6) and in SN4741(Fig. 7).

We have tried to find possible explanations for the discrepancy between our results and those of Wang et al [12]. We found differences in the protein extraction buffer. We routinely use SDS-Laemmli loading buffer and they use a different buffer to prepare the total cell lysates before missing with SDS-loading buffer, protein extraction under their conditions gave in our hands similar results to those presented here. Another difference is the anti-DJ-1 antibody, they use Abcam, ab76008. We bought that antibody and in our hands gave similar results to Abcam ab18257 used in the present work and we also obtained the same results using our in-house produced polyclonal anti-DJ-1 antibody [10].

Our results are in agreement with previous reported results on the stability of DJ-1 from typical biochemical studies [9] [11] [10]. Furthermore, quantitative proteomic studies using SILAC pulse-chase experiments and MS analysis also show that DJ-1 protein has a very long half-life (t1/2). For example; DJ-1 has a t1/2=187h in mouse NIH3T3 cells [19], the values in HeLa and C2C12 are t1/2=59.2h and t1/2= 88h, respectively [20]. In vivo studies by deuterium labelling and MS also show that the t1/2 of DJ-1 in heart from mice varies from 199 to 299h depending of the mouse strain under study [21]. Finally, recent studies by proteomic MS of the degradation of “young” (newly synthesized) and “old” (steady state) proteins in NIH3T3 report that DJ-1 degradation follows an exponential decay, exponential 1-state t1/2 = 150.26h and steady-state 2- state model with a t1/2 = 217.1h and also exponential decay is observed in human RPE-1 cells with a t1/2 >300h [22]. In all cases DJ-1 protein has a very long half-life, several days. These facts, together with the ubiquitous expression of DJ-1, are in agreement with the proposal to use DJ-1 peptides as reference for quantification of proteomic studies, being even better that commonly used peptides from house keeping proteins [23]. With a half-life of several days, even for the newly synthesized DJ-1 protein [22], the study of the pathway responsible for the degradation of DJ-1 is not easy to approach experimentally by applying treatments that can stimulate or inhibit its degradation. In particular, it could be that longer serum fasting times (> 24h) may be required to observe a significant change in DJ-1 protein levels. Unfortunately those experiments are not possible, because the cells (MEFs, N2a and B-LCL) used in this work do not survive longer fasts (36-48h). Using HeLa cells, that are more resistant, no change in DJ-1 protein levels were observed even after 48h of serum starvation.

In conclusion, DJ-1 protein has a long half-life in the cell and we have not found any change in DJ-1 protein levels by inactivation of LAMP-2 gene expression, inhibition of lysosomal function or activation of the CMA pathway by serum starvation.

Accordingly, we have been unable to reproduce the observations reported by Wang et al [12] on the role of CMA in DJ-1 protein turn-over.

## Acknowledgments

Special thanks to Drs. Judith Blanz and Paul Saftig from Institute of Biochemistry, Christian-Albrechts-Universität zu Kiel, Olshausenstrasse 40, D-24098 Kiel, Germany for providing us the MEF and N2a cell lines used in the present work and for critical reading of the manuscript. We thank José Luis Zugaza from Department of Genetics, Physical Anthropology and Animal Genetics, Faculty of Science and Technology, University of the Basque Country, UPV/EHU and Achucarro Basque Center for Neuroscience, Bilbao, Spain for providing us with the SN4741 cell line. This work was supported by grants from MINECO SAF-2012-34556 and CIBERNED to JGC.

